# Small variant benchmark from a complete assembly of X and Y chromosomes

**DOI:** 10.1101/2023.10.31.564997

**Authors:** Justin Wagner, Nathan D. Olson, Jennifer McDaniel, Lindsay Harris, Brendan J. Pinto, David Jáspez, Adrián Muñoz-Barrera, Luis A. Rubio-Rodríguez, José M. Lorenzo-Salazar, Carlos Flores, Sayed Mohammad Ebrahim Sahraeian, Giuseppe Narzisi, Marta Byrska-Bishop, Uday S Evani, Chunlin Xiao, Juniper A. Lake, Peter Fontana, Craig Greenberg, Donald Freed, Mohammed Faizal Eeman Mootor, Paul C. Boutros, Lisa Murray, Kishwar Shafin, Andrew Carroll, Fritz J Sedlazeck, Melissa Wilson, Justin M. Zook

**Affiliations:** National Institute of Standards and Technology, Material Measurement Laboratory, 100 Bureau Dr., Gaithersburg, MD 20899, USA; Center for Evolution & Medicine, Arizona State University, Tempe, AZ 85281 USA -- Department of Zoology, Milwaukee Public Museum, Milwaukee, WI 53233 USA; Genomics Division, Instituto Tecnológico y de Energías Renovables (ITER), Granadilla de Abona, Spain; CIBER de Enfermedades Respiratorias (CIBERES), Instituto de Salud Carlos III, Madrid, Spain; Research Unit, Hospital Universitario Ntra. Sra. de Candelaria, Santa Cruz de Tenerife, Spain; Facultad de Ciencias de la Salud, Universidad Fernando de Pessoa Canarias, Las Palmas de Gran Canaria, Spain; Roche Sequencing Solutions, Santa Clara, CA, USA; New York Genome Center, NewYork, NY 10013, USA; National Center for Biotechnology Information, National Library of Medicine, National Institutes of Health, Bethesda, MD 20894 USA; Pacific Biosciences, Menlo Park, CA, USA; National Institute of Standards and Technology, Information Technology Laboratory, 100 Bureau Dr. Mailstop 8940, Gaithersburg, MD 20899, USA; Sentieon Inc. San Jose, CA, USA; Department of Human Genetics, University of California Los Angeles, Los Angeles, CA, USA; Illumina, San Diego, CA, USA; Google Inc, 1600 Amphitheatre Pkwy, Mountain View, CA, USA; Baylor College of Medicine Human Genome Sequencing Center, Houston, TX, USA; Center for Evolution & Medicine and School of Life Sciences, Arizona State University, Tempe, AZ 85281 USA

## Abstract

The sex chromosomes contain complex, important genes impacting medical phenotypes, but differ from the autosomes in their ploidy and large repetitive regions. To evaluate variant detection on chromosomes X and Y, we created an 111,725 variant benchmark for the Genome in a Bottle HG002 reference material. We show how complete assemblies can expand benchmarks to difficult regions, but highlight remaining challenges benchmarking complex gene conversions, copy number variable gene arrays, and human satellites.

## Main Text

The complete human karyotype includes 22 pairs of autosomes and two sex chromosomes (X and Y). The unique biology of the X and Y chromosomes makes their analysis more difficult than the autosomes in some ways. Indeed, the X and Y chromosomes contain many medically relevant genes, as well as very challenging repetitive regions.^1–5^ Chromosomes X and Y mostly have distinct sequences, but two pseudoautosomal regions (PARs) experience crossover events similar to autosomes, and the recently X transposed region (XTR) retains relatively high sequence identity between X and Y.^6^ Benchmark sets from well-characterized samples are important for understanding variant call accuracy. Previous Genome in a Bottle Consortium (GIAB) benchmarks excluded the X and Y chromosomes due to their mostly-hemizygous (i.e., haploid) nature in half the population,^7,8^ which requires customized variant calling methods.^9^ However, recently, GIAB has developed approaches to form variant benchmarks by aligning long-read assemblies to the reference,^10,11^ enabling the generation of benchmarks for more challenging regions and variants. Here, we create benchmarks that include challenging variants and regions by using complete, polished *de novo* assemblies of the X and Y chromosomes in the GIAB Personal Genome Project^12^ Ashkenazi Jewish son HG002 from the Telomere-to-Telomere (T2T) Consortium.^1,13^ We also pilot a new systematic approach to evaluate benchmarks to exclude problematic regions and ensure the final benchmark reliably identifies errors in a variety of genome contexts.

Our new benchmark includes many but not all challenging regions. To create a small variant benchmark for SNVs and small indels less than 50 bp in size, we aligned the T2T assembly of HG002’s chromosomes X and Y to GRCh38 and called variants with dipcall.^14^ We then excluded regions where assembly-based small variant calling or benchmarking tools were unreliable, such as regions containing structural variants, as detailed in Online Methods.

The resulting benchmark contains substantially more challenging variants and regions than previous benchmarks. It includes 94% of chromosome X and 63% of chromosome Y, after excluding the 0.7% of chromosome X and 53% of chromosome Y missing from GRCh38. Of 294 medically relevant genes on these chromosomes,^8^ 270 are >90% included; of which 251 are >99% included in the small variant benchmark regions. The benchmark includes some challenging regions like 68% of segmental duplications, and 99% of the XTR.

Although the new benchmark excludes the most challenging regions, it correctly identifies many errors in challenging regions, including variants missed by short- and long-read mapping-based approaches. Table 1 describes how older short-read and long-read variant calls from the 2020 precisionFDA Challenge^15,16^ perform substantially worse against the assembly-based autosomal Challenging Medically Relevant Gene (CMRG)^8^ and XY benchmarks relative to the v4.2.1 benchmark,^7^ which used mapping-based approaches except in the major histocompatibility complex region. This drop in performance is particularly evident for SNVs in segmental duplications, large indels, and indels in tandem repeats and homopolymers. The lower F1 score for insertions > 15 bp seen against the XY benchmark, especially with long reads, can be attributed to a higher rate of genotype errors. The benchmark regions also have substantially lower variant density on Y (170/Mbp) compared to X (746/Mbp), as expected from previous work.^17^

**Table 1:** F1 scores for short-read (Illumina with DeepVariant) and long-read callsets (PacBio HiFi with DeepVariant) from 2020 precisionFDA challenge against three recent GIAB benchmarks: the mapping-based v4.2.1 small variant benchmark for autosomes, the small variant benchmark for 273 challenging medically-relevant autosomal genes (CMRG) in autosomes, and the new XY benchmark in this work. These do not reflect current accuracy, but exemplify how performance metrics can depend substantially on the difficulty of variants and regions included in the benchmark.

| Variant Type | Region | Short read (2020) |  |  | Long read (2020) |  |  |
| --- | --- | --- | --- | --- | --- | --- | --- |
|  |  | v4.2.1 | CMRG | XY | v4.2.1 | CMRG | XY |
| SNV | All benchmark | 0.997 | 0.977 | *0.899* | 1.000 | 0.981 | *0.932* |
| SNV | Segmental duplications | 0.951 | 0.835 | *0.600* | 0.991 | 0.893 | *0.785* |
| INDEL | All benchmark | 0.997 | 0.963 | *0.815* | 0.996 | 0.967 | *0.738* |
| INDEL | TRs | 0.993 | 0.915 | *0.721* | 0.997 | 0.955 | *0.645* |
| INDEL | Homopolymers longer than 11bp | 0.998 | 0.972 | *0.789* | 0.990 | 0.959 | *0.677* |
| Insertions >15bp | All benchmark | 0.960 | 0.821 | *0.538* | 0.997 | 0.919 | *0.505* |

By using alignments of the complete assembly to the GRCh38 reference, we included some gene conversion-like events in the benchmark, where in HG002 the sequence of one GRCh38 region is replaced by the sequence of another similar GRCh38 region. For example, the genes *MAGEA6* and *MAGEA3* are swapped and inverted in HG002 relative to GRCh38, as described previously^18^ (Figure 1). The gene *CSAG2* is in the same segmental duplication in GRCh38, and in HG002 the region more closely matches CSAG3. Unlike MAGEA3 and MAGEA6, which are swapped, HG002 also still has *CSAG3* in its location on GRCh38, so it is more similar to a stereotypical intrachromosomal gene conversion event. Even further towards the middle of these segmental duplications, *MAGEA2* and *MAGEA2B* are identical in GRCh38, and are also almost identical to each other in HG002, but their sequences in HG002 are diverged from GRCh38. In the middle of this pair of segmental duplications, *CSAG1* and *MAGEA12* have some SNVs but do not appear inverted or converted.

**Figure 1:**
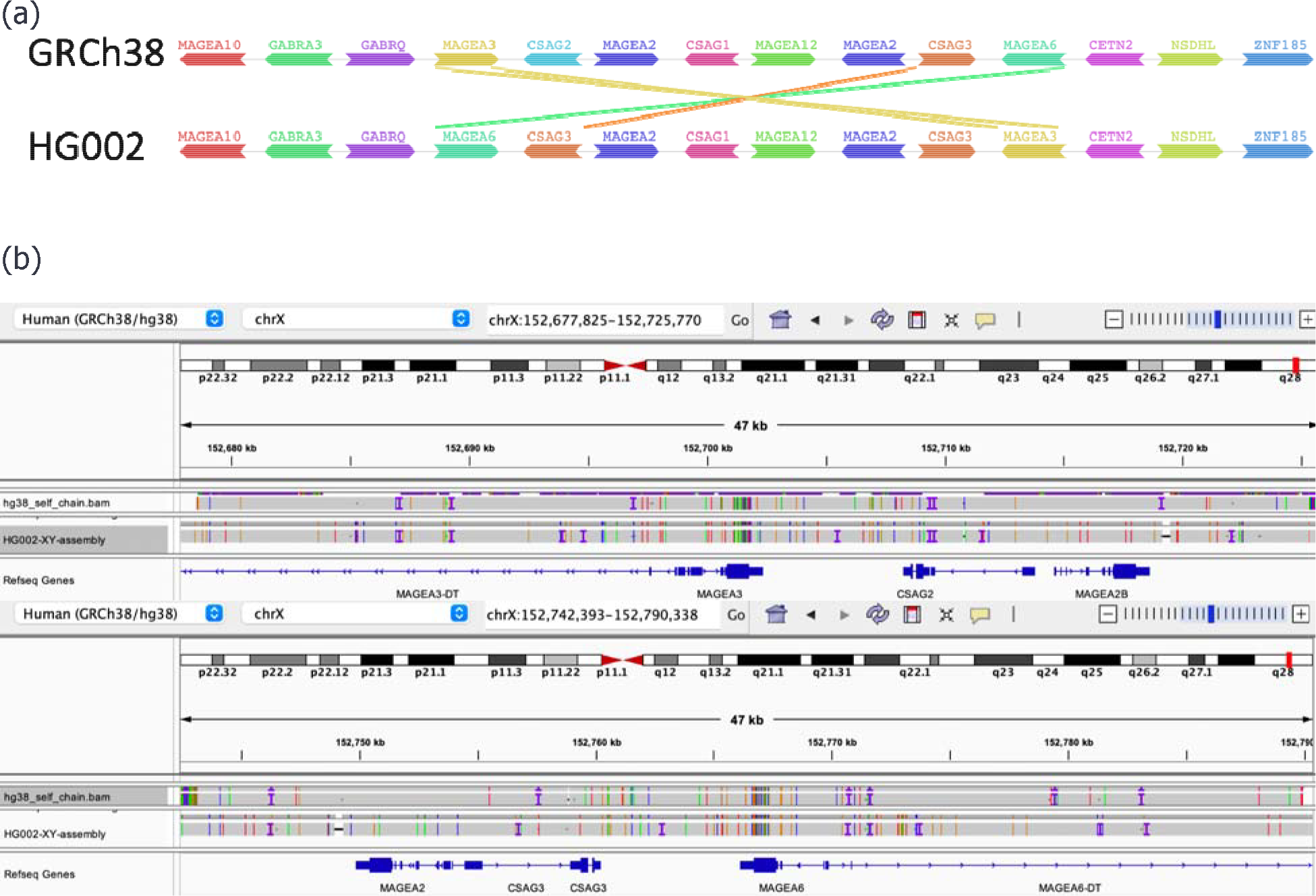
Visualization of gene conversion events included in the benchmark. (a) Visualization of genes in a region containing an inverted segmental duplication pair using the pangene tool (more haplotypes can be seen at http://pangene.liheng.org/view?graph=human98-r183a&gene=MAGEA3&step=3&ori=). (b) IGV visualization showing that many of the variants in *MAGEA3, CSAG2*, and *MAGEA6* match the variants in the self-chain alignment of the segmental duplication, indicating a gene conversion-like event.

Beyond gene conversion-like events, the benchmark includes other challenging variants and regions. For example, Supplementary Figure 1 shows an example of complex variants in a tandem repeat in the PAR, where the assembly resolves multiple phased variants on each haplotype within the repeat, which can be challenging for mapping-based approaches. Another challenging region for variant calling in the benchmark is a 856 kbp region on the Y chromosome with a high density of false positives (FPs) due to extra divergent copies of the sequence in HG002 but not GRCh38, shown in Supplementary Figure 2. The assembly resolves the correct sequence in this region, whereas mapping-based methods contain many FPs due to mismapped reads.

We excluded other challenging regions due to errors in the HG002 assembly and ambiguous assembly-assembly alignments around large, complex SVs. For example, we excluded a large known inversion error in the HG002 assembly on chrY. We also excluded the TSPY2 gene, which undergoes a complex interchromosomal gene conversion, as well as the TSPY gene array, which has 46 copies in HG002 versus 9 copies in GRCh38. We detail these challenges in Supplementary Note 3, highlighting a remaining gap in complete variant benchmarks,

These results demonstrate the ongoing need for better benchmarking tools for tandem repeats, large duplications, and complex SVs. Even where assemblies are correct, there is a need for improved assembly-assembly alignment methods and standards for representing and comparing variants in regions that do not have 1:1 alignment between assembly and reference.^19^ The clearest example of this is *TSPY2*, excluded in this current benchmark with plans to revisit this region as part of the HG002 T2T diploid assembly project. Also, a small number of structural errors remain in these complete X and Y assemblies, such as the inversion on chromosome Y that is polymorphic in the population as identified by Ref^1^. Additionally, there are still many errors in homopolymers and dinucleotide tandem repeats in current assemblies which are not completely captured by kmer-based methods typically used to assess assembly error rates. A final consideration is to exclude mosaic variants, which we are in the process of identifying and characterizing. We acknowledge an additional limitation of the current HG002 XY v1.0 benchmark that it may be biased towards HiFi and Element as a result of assembly input and regions excluded. The HG002 T2T Q100 effort (https://github.com/marbl/HG002) to polish diploid assemblies will fill this gap and enable inclusion of homopolymers in the benchmark that are noisy in HiFi. This manuscript provides an important initial benchmark from two T2T chromosomes, and highlights the strengths of this approach as well as remaining challenges for T2T benchmarks for the whole genome.

## Online Methods

### Benchmark Generation

Based on previous GIAB work,^8^ we included only loci with exactly one contig aligning from each haplotype (except in the X and Y non-PAR), which is the dip.bed file output from dipcall. Furthermore, we excluded structural variants at least 50 bp in size and associated repeats, large repeats (segmental duplications, tandem repeats longer than 10 kbp, and satellites) that have any breaks in the assembly-assembly alignment, and regions around gaps in GRCh38.^20^ This was implemented as a snakemake pipeline (https://github.com/nate-d-olson/defrabb). The version of the defrabb repo used to generate the XY benchmark was https://github.com/nate-d-olson/defrabb/tree/b0f08b6b0514555570e8f90fa51b0a86a3c904da. The config files used for the defrabb run, include input files for the repeats, are at https://github.com/nate-d-olson/defrabb/blob/b0f08b6b0514555570e8f90fa51b0a86a3c904da/config/analyses_20230315_v0.011-HG002XY.tsv and https://github.com/nate-d-olson/defrabb/blob/b0f08b6b0514555570e8f90fa51b0a86a3c904da/config/resources.yml.

We refined the benchmark using an active evaluation approach (https://github.com/usnistgov/active-evaluation) as we detail in the Supplementary Note 1. Based on curation of 12 callsets compared against the draft benchmark (see “Visualizing and curating variants to understand errors” of Olson *et al*.^21^), we further excluded homopolymers longer than 30 bp, tandem repeats discordant with a different HG002 assembly, homopolymers discordant with Element avidity-based sequencing variant calls, as well as regions with ambiguous or incorrect assembled sequence or assembly-assembly alignments (Supplementary Note 2).

### Evaluation

We evaluated the benchmark to test our criteria for Reference Materials to be fit-for-purpose, which in this case is the reliable identification of errors in diverse variant callsets.^21^ In this work, we piloted a method for evaluating machine learning systems called Active Evaluation that takes advantage of stratifying a dataset and estimates a confidence interval for a system’s performance. We defined 15 stratifications - as shown in Supplementary Figure 3 - based on homopolymer length, difficult-to-map regions, and pseudoautosomal regions, as well as SNVs vs indels and putative false positives versus false negatives (Supplementary Note 1 and Supplementary Figure 4). We additionally performed long range PCR followed by Sanger sequencing on a subset of variants in challenging genes in chromosomes X and Y as a means of orthogonal validation of the benchmark variants (Supplementary Note 1 and Supplementary Table 2).

## Data availability

The benchmark vcf and bed, as well as supporting files, are available at https://ftp-trace.ncbi.nlm.nih.gov/ReferenceSamples/giab/release/AshkenazimTrio/HG002_NA24385_son/chrXY_v1.0/

## Competing interests

JAL is an employee of PacBio. DF is an employee of Sentieon, Inc., and holds stock options as part of the standard compensation package. PCB sits on the Scientific Advisory Boards of Intersect Diagnostics Inc., Sage Bionetworks and BioSymetrics Inc. LM is an employee and shareholder of Illumina Inc. KS and AC are employees of Google LLC and own Alphabet stock as part of the standard compensation package. FJS has support from ONT, Illumina, Pacbio and Genentech.

## Acknowledgements

Certain commercial equipment, instruments, or materials are identified to specify adequately experimental conditions or reported results. Such identification does not imply recommendation or endorsement by the National Institute of Standards and Technology, nor does it imply that the equipment, instruments, or materials identified are necessarily the best available for the purpose.

## Notes

https://ftp-trace.ncbi.nlm.nih.gov/ReferenceSamples/giab/release/AshkenazimTrio/HG002_NA24385_son/chrXY_v1.0/

### Summary of Updates

Shortening and minor updates to text

