## Supplementary Notes and Figures for "Small variant benchmark from a complete assembly of X and Y chromosomes"

### *Supplementary Note 1: Benchmark Set Evaluation*

We defined 15 stratifications - as shown in Supplementary Figure 3 - based on homopolymer length, difficult-to-map regions, and pseudoautosomal regions, as well as SNVs *vs* indels and putative false positives versus false negatives. We then produced a human curation for sampled variant calls using binary labels denoting whether the benchmark is correct for that variant. We stratified and manually curated putative false positives and false negatives for 12 callsets against a draft XY benchmark for HG002.

The results of the evaluation and manual curation are shown in Supplementary Figure 4 and Supp Table 1. We found that, overall, the benchmark reliably identifies errors across stratifications and callsets. One notable exception was for indel FPs and FNs in an Element callset, where most of the putative FPs and FNs were errors in the benchmark in long homopolymers. Element's avidity sequencing has been demonstrated to have high accuracy in homopolymers.[22] Therefore, during our evaluation effort, we refined the benchmark by excluding additional region types, as described above. We also excluded all regions containing variants identified as incorrect or 'unsure' during manual curation.

We performed manual curation of a subset of putative False Positive and False Negative calls resulting from a comparison of the draft benchmark against a variety of callsets from different technologies. This effort was similar to the evaluation process of the v4.2.1[7] and CMRG[8] GIAB benchmarks. We updated this approach using a more focused sampling of different subsets or evaluation strata of the comparison result. We then used an Active Evaluation approach (https://github.com/usnistgov/active-evaluation) to estimate a score indicating how often the benchmark matched the manual curation result as well as confidence intervals for each score. In using this approach, the overall goal is that the benchmark has a score of greater than 0.5 - or match the manual curation result in more than 50% of instances with 96% confidence intervals entirely above 0.5 indicating that the true estimate of the benchmark matching a manual curation result lies above 50% of instances. Overall, as seen in Supplementary Table 1 across all callsets the lower confidence interval is above 0.5 and in many of the callsets is significantly higher. To see this, look at the column sys_conf_lower, which gives the lower bound of the confidence intervals. As the lowest value is 0.553 (rounded to 3 decimal places), this means that all the system 95% confidences are above 0.50, meaning that if we were to curate the entire population it is likely that the benchmark agrees with the curation more than 50% of the time. This provides users evidence and well-defined confidence intervals that the benchmark is fit for purpose to reliably identify errors in other callsets.

We also performed long range PCR followed by Sanger sequencing on a subset of challenging variants in chromosomes X and Y as a means of orthogonal validation of the benchmark variants. We confirmed a total of 181 variants in segmental duplications in 10 genes: *ARHGAP6, CLIC2, CSAG1, F8, IKBKG, NXF5, OPN1LW, OPN1MW, SAGE1, SLC6A8, SLC6A14, TMLHE.* Only one variant appeared contradicted by Sanger, where the assembly was clearly supported by long reads and the reason for the different Sanger result was unclear. The variant confirmation results and experimental conditions are detailed in Supp Table 2.

### *Supplementary Note 2: Calculations to create the benchmark*

The description of calculations used to create the final v1.0 benchmark are in this notebook: [https://github.com/jmcdani/giab-chrXY-benchmark/blob/main/scripts/chrXY_benchmark_exclusions.ipynb](https://github.com/jmcdani/giab-chrXY-benchmark/blob/main/scripts/chrXY_benchmark_exclusions.ipynb)

### *Supplementary Note 3: Remaining challenging regions excluded from the benchmark*

We highlight two examples of challenging segmental duplications on chromosome Y that we excluded from the benchmark. The first region is a small known inversion error in the HG002 assembly[1] flanked by a pair of very large segmental duplications (chrY:17,455,804-17,951,260 and chrY:17,954,718-18,450,201). The intervening sequence between these regions is incorrectly inverted and the flanking segmental duplications also contain some small errors. Thus, we excluded the entire region including the segmental duplications (chrY:17,455,804-18,450,201).

The second set of regions excluded involves two different assembly-assembly alignment challenges posed by the TSPY gene family. The TSPY gene family is highly polymorphic in copy number[23] and has been implicated in risk of infertility and cancer.[24,25] The first challenge is that segmental duplications including the *TSPY2* gene were found to be swapped with their homologous sequences about 4 Mbp downstream in HG002 and most individuals relative to GRCh38.[23] We found that standard assembly-based variant calling methods such as dipcall, used in our work, tend to align the assembly contiguously rather than breaking the alignment and aligning *TSPY2* in HG002 to *TSPY2* in GRCh38. This results in variant calls where *TSPY2* is mostly deleted at its location in GRCh38 and inserted in its location in HG002. It is not standardized whether the variants should be called in this way or whether *TSPY2* in HG002 should be aligned to *TSPY2* in GRCh38, resulting in large translocations and smaller variants. Therefore, we chose to exclude these regions from the current benchmark bed file. Since there are multiple segmental duplication pairs annotated in this region of GRCh38 that appear to be swapped in HG002, we specifically excluded the segmental duplication pairs in these regions: chrY:6,234,812-6,532,742 and chrY:9,628,425-9,919,592. In addition to TSPY2 moving about 4 Mbp, the nearby gene *TTTY22* as well as *RBMY2NP, RMBY2GP* and some other TTTY paralogs also swap positions between the segmental duplications chrY:6,234,812-6,532,742 and chrY:9,628,425-9,919,592. Interestingly, all of these genes are annotated differently on T2T-Y by CAT+Liftoff and RefSeq. Where RefSeq seems to match gene sequences, CAT+Liftoff tries to match positions even if the gene sequences differ.

In addition to the *TSPY2* challenge, the TSPY gene array is expanded substantially in HG002 relative to GRCh38, with 46 copies in HG002 versus 9 copies in GRCh38, which has a 40 kb gap (see "Ampliconic genes in composite repeats" in Ref[1]). In this case, no standards exist for whether this should be represented as one or more very large insertions alongside smaller variants, as a copy number variant, as a tandem repeat expansion, or some alternate representation. For these reasons, we excluded the TSPY gene array from the benchmark.

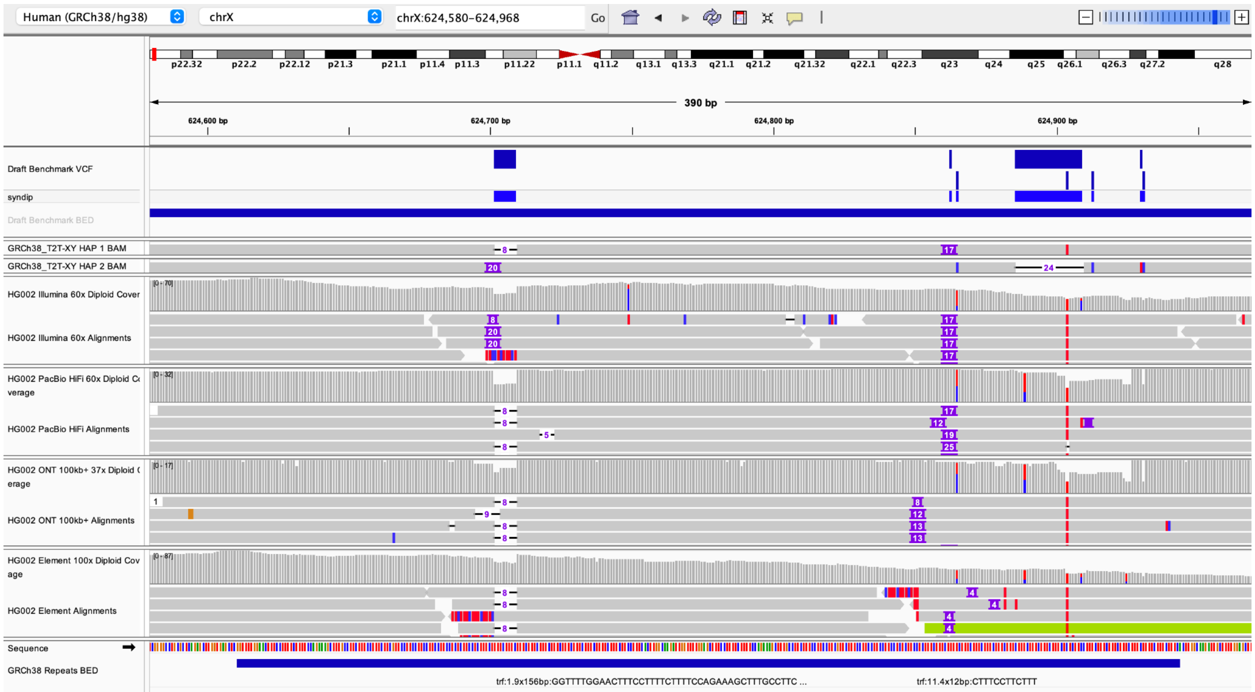


**Supplementary Figure 1**: Complex variants in a long tandem repeat in the PAR. In this region, the assemblies used for the benchmark accurately resolve horizontally complex variants, with multiple variants in each haplotype, resulting in phased variant calls that can be used for benchmarking.

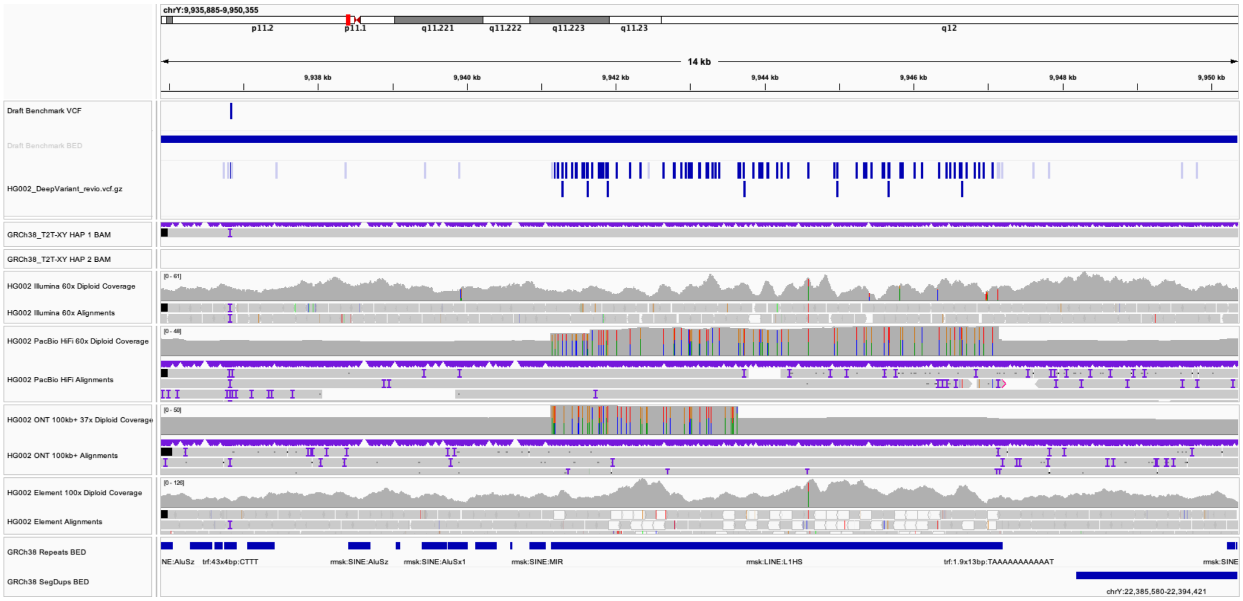


**Supplementary Figure 2**: Region with many false positives in mapping-based methods due to additional divergent copies of sequence in HG002 not in GRCh38. The assembly resolves the correct sequence in this region (chrY:10744000-11600000), with only one variant relative to GRCh38. Mapping-based approaches can result in many false positives due to differences in reads mis-mapping from the additional copies of this sequence in HG002.

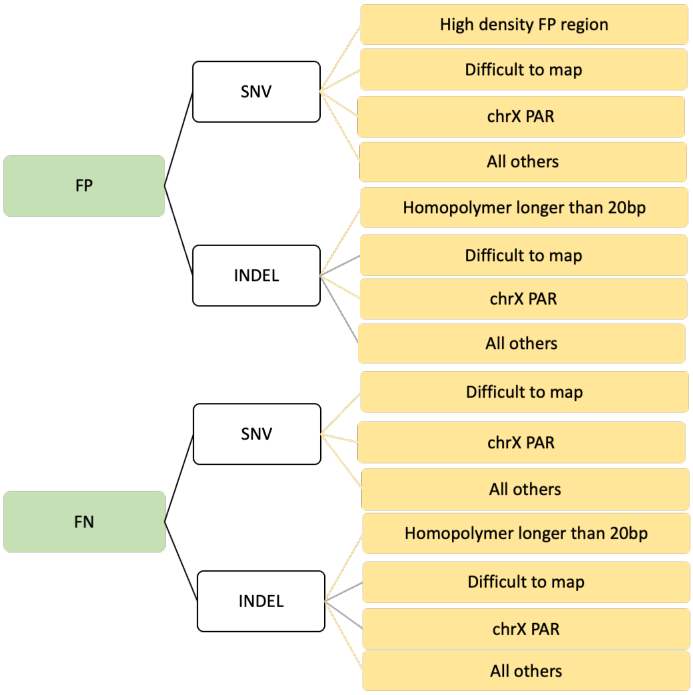


**Supplementary Figure 3**: Strata Descriptions used during evaluation. Putative False Positive and False Negative SNVs and INDELs were categorized into different genomic regions. These include the High Density FP region shown in Supplementary Figure 2, difficult to map regions, the PAR of chrX, homopolymers longer than 20bp, and all others outside these regions.

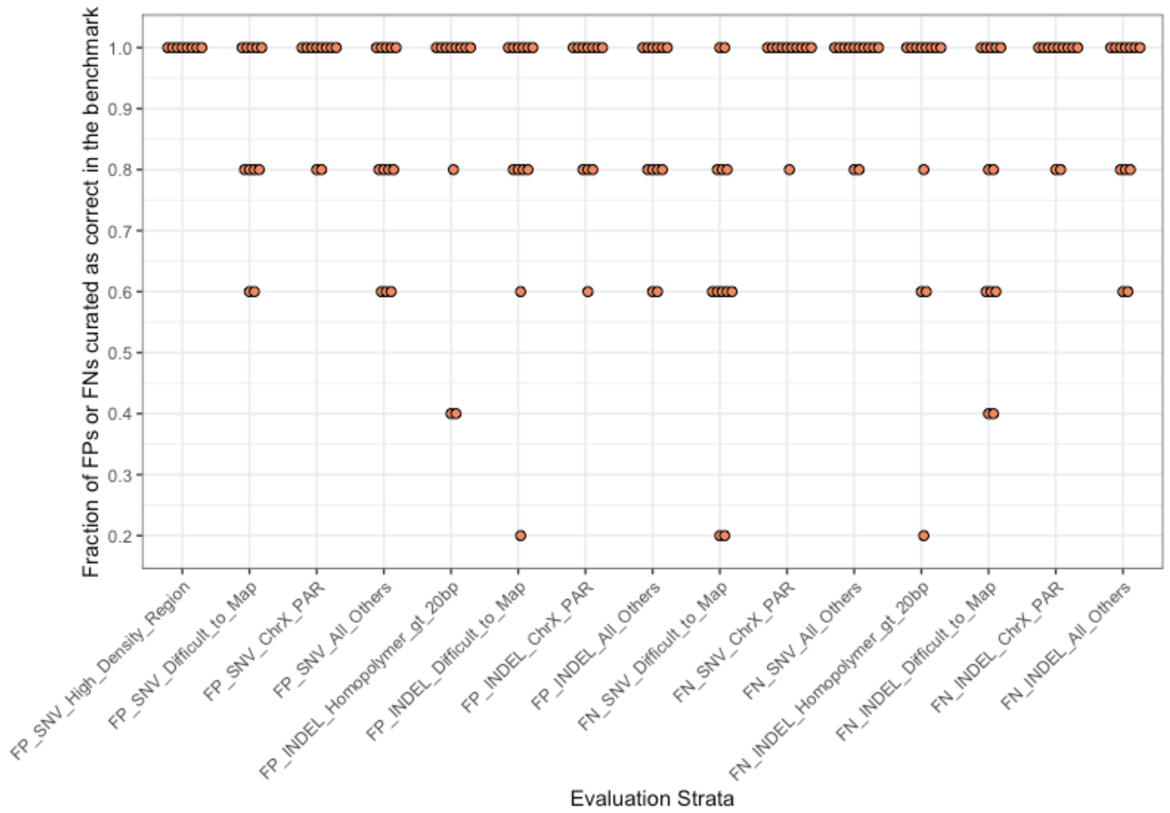


**Supplementary Figure 4**: Evaluation of benchmark against a variety of sequencing technologies and variant calling methods. Each point represents curation results for each variant callset against draft benchmarks with the points below 0.5 as support for excluding regions from the v1.0 benchmark. Additionally, all variants curated as unsure or incorrect in the benchmark were excluded from the v1.0 benchmark regions.
